# Performance of neural network basecalling tools for Oxford Nanopore sequencing

**DOI:** 10.1101/543439

**Authors:** Ryan R. Wick, Louise M. Judd, Kathryn E. Holt

**Affiliations:** Department of Infectious Diseases, Central Clinical School, Monash University, Melbourne, Victoria 3004, Australia; London School of Hygiene & Tropical Medicine, London WC1E 7HT, UK

## Abstract

**Background:** Basecalling, the computational process of translating raw electrical signal to nucleotide sequence, is of critical importance to the sequencing platforms produced by Oxford Nanopore Technologies (ONT). Here we examine the performance of different basecalling tools, looking at accuracy at the level of bases within individual reads and at majority-rules consensus basecalls in an assembly. We also investigate some additional aspects of basecalling: training using a taxon-specific dataset, using a larger neural network model and improving consensus basecalls in an assembly by additional signal-level analysis with Nanopolish.

**Results:** Training basecallers on taxon-specific data results in a significant boost in consensus accuracy, mostly due to the reduction of errors in methylation motifs. A larger neural network is able to improve both read and consensus accuracy, but at a cost to speed. Improving consensus sequences (‘polishing’) with Nanopolish somewhat negates the accuracy differences in basecallers, but pre-polish accuracy does have an effect on post-polish accuracy.

**Conclusions:** Basecalling accuracy has seen significant improvements over the last two years. The current version of ONT’s Guppy basecaller performs well overall, with good accuracy and fast performance. If higher accuracy is required, users should consider producing a custom model using a larger neural network and/or training data from the same species.

## Background

Oxford Nanopore Technologies (ONT) long read sequencing is based on the following concept: pass a single strand of DNA through a membrane via a nanopore and apply a voltage difference across the membrane. The nucleotides present in the pore will affect the pore’s electrical resistance, so current measurements over time can indicate the sequence of DNA bases passing through the pore. This electrical current signal (a.k.a. the ‘squiggle’ due to its appearance when plotted) is the raw data gathered by an ONT sequencer. Basecalling for ONT devices is the process of translating this raw signal into a DNA sequence. It is not a trivial task as the electrical signals come from single molecules, making for noisy and stochastic data. Furthermore, the electrical resistance of a pore is determined by the bases present within multiple nucleotides that reside in the pore’s narrowest point (approximately five nucleotides for the R9.4 pore), yielding a large number of possible states: 4^5^=1024 for a standard four-base model. When modified bases are present, e.g. 5-methylcytosine, the number of possible states can grow even higher: 5^5^=3125. This makes basecalling of ONT device signals a challenging machine learning problem and a key factor determining the quality and usability of ONT sequencing.

Basecalling is an active field, with both ONT and independent researchers developing methods. Modern basecallers all use neural networks, and these networks must be trained using real data. The performance of any particular basecaller is therefore influenced by the data used to train its model. This is especially relevant when basecalling native (not PCR-amplified) DNA, which can contain base modifications. A basecaller’s performance in such a case may depend on whether the modifications and their sequence motifs were represented in its training set.

Basecalling accuracy can be assessed at the read level (read accuracy) or in terms of accuracy of the consensus sequence (consensus accuracy). Read accuracy measures the sequence identity of individual basecalled reads relative to a trusted reference. Consensus accuracy measures the identity of a consensus sequence constructed from multiple overlapping reads originating from the same genomic location. Consensus accuracy generally improves with increased read depth, e.g. a consensus built from 10 reads is likely to be less accurate than one built from 100 reads.

While read and consensus accuracy may be correlated, this relationship is not guaranteed. I.e. more accurate reads do not necessarily produce a more accurate consensus. Random read errors are unlikely to appear in the consensus, as they occur in the minority of reads at their locus. Systematic errors that occur in many reads can however appear in the consensus. Low accuracy reads can therefore produce a perfect consensus sequence, provided their errors are random and the read depth is sufficiently large. Conversely, high accuracy reads can create an imperfect consensus regardless of the read depth, if they contain systematic errors.

Consensus accuracy is usually the main concern for applications with high read depth, such as genome assembly. For other applications, particularly those with low read depths, read accuracy is important. For example, clinical metagenomics may rely on data from a very small number of non-human reads^1^, and inaccurate reads could make it harder to identify and characterise pathogens.

ONT have released multiple pore types during their history, but the R9.4 pore (and its minor revision R9.4.1) has been available for longest: from October 2016 to the present. Its release also corresponds to approximately when command-line basecallers became available – before that users needed to basecall with ONT’s cloud-based Metrichor service. This study aims to quantify the performance of various basecalling tools developed for ONT’s R9.4 pore and to explore the impact of model training on basecalling accuracy. It may provide guidance to those wishing to get the most out of ONT sequencing signals, and in particular could help readers to decide whether recent progress warrants re-basecalling older signal data with a newer basecaller or custom-trained model.

In this study, we tested four basecalling programs developed by ONT – Albacore, Guppy, Scrappie and Flappie – and ran all available versions compatible with R9.4 reads. Albacore is a general-purpose basecaller that runs on CPUs. Guppy is similar to Albacore but can use GPUs for improved basecalling speed. While the two basecallers have coexisted for about a year, ONT has discontinued development on Albacore in favour of the more performant Guppy. Both Albacore and Guppy are only available to ONT customers via their community site (community.nanoporetech.com). Scrappie (github.com/nanoporetech/scrappie) is an open-source basecaller which ONT describes as a ‘technology demonstrator’. It has often been the first of ONT’s basecallers to try new approaches, with successes later being incorporated into Albacore and Guppy. Scrappie is really two basecallers in one: Scrappie events, which carries out an event-segmentation step prior to basecalling with its neural network, and Scrappie raw, which basecalls directly from raw signal. We excluded some older versions of Scrappie events which rely on events first being defined by another program, as this requirement makes it not a standalone basecaller. Flappie (github.com/nanoporetech/flappie) has recently replaced Scrappie and uses a CTC decoder to assign bases^2^. We also tested Chiron (github.com/haotianteng/Chiron), a third-party basecaller still under development that uses a deeper neural network than ONT’s basecallers^3^. We excluded older basecallers no longer under development, such as Nanonet, DeepNano^4^ and basecRAWller^5^.

## Results and discussion

### Default model performance

Our primary benchmarking read set for assessing basecaller performance consisted of 15 154 whole genome sequencing reads from an isolate of *Klebsiella pneumoniae* (see Methods). These reads ranged from 22-134kbp in length (N50=37 kbp) and were ∼550Mbp in total size, equating to 100× read depth over the 5.5 Mbp *K. pneumoniae* chromosome.

Albacore’s history contained two major developments which resulted in distinct improvements in both read and consensus accuracy: in April 2017 (version 1.0.1) and August 2017 (version 2.0.1) (Fig 1). The first was the addition of a transducer to the basecaller^6^, which allowed for better homopolymer calls (see error profile details below). The second was the switch to raw basecalling, where sequence is called directly from raw signal without an event-segmentation step. After August 2017, Albacore’s performance remained fairly constant with subsequent releases, achieving read accuracy of Q9.2 and consensus accuracy of Q21.9 with its final version (v2.3.4).

**Fig 1.**
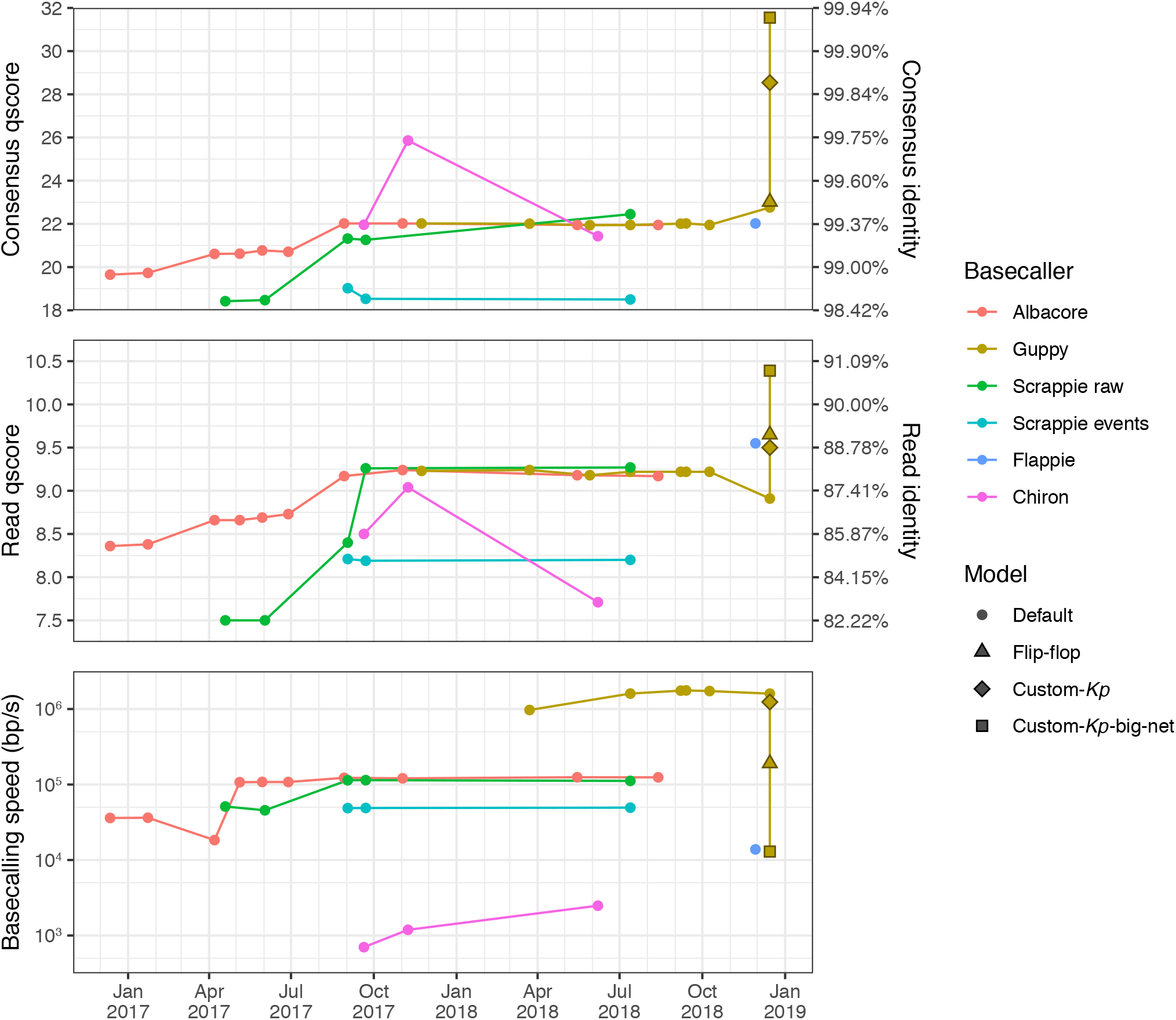
Read accuracy, consensus accuracy and speed performance for each basecaller version, plotted against the release date (version numbers specified in File S1). Accuracies are expressed as qscores (also known as Phred quality scores) on a logarithmic scale where Q10=90%, Q20=99%, Q30=99.9%, etc. Each basecaller was run using its default model, except for Guppy v2.2.3 which was also run with its included flip-flop model and our two custom-trained models.

Guppy was publicly released in late 2017 (v0.3.0) and its accuracy stayed relatively constant and similar to that of Albacore for most of its version history (up to v1.8.5 in October 2018). The last tested version of Guppy (v2.2.3, released January 2019) performed worse on read accuracy (Q8.9) but better on consensus accuracy (Q22.8) using its default model. This version also comes with an optional ‘flip-flop’ model which has similar consensus accuracy (Q23.0) but much better read accuracy (Q9.7). However, the flip-flop model’s accuracy comes at the cost of speed performance – it is considerably slower than the default Guppy model, likely due to its larger size and complexity (Fig S2).

Scrappie events was the worst-performing basecaller tested, likely due to its use of an outmoded event-segmentation pre-processing step. Scrappie raw performed better, and the latest version (v1.4.1) performs similarly to Albacore (Q9.3 read accuracy and Q22.4 consensus accuracy). Scrappie’s successor Flappie (released November 2018) showed an improvement in read accuracy (Q9.6) but not consensus accuracy (Q22.0).

Relative to ONT’s basecallers, Chiron performed poorly on read accuracy (Fig 1). Chiron v0.3 had the highest consensus accuracy (Q25.9) of all tested basecallers using their default models, but the latest version (v0.4.2) did not perform as well on our benchmarking set (Q7.7 read accuracy and Q21.4 consensus accuracy). This variation in accuracy for different versions of Chiron is explained by the taxonomy of the training data used to produce their default models (*E. coli* vs human, see Consensus error profiles).

While Albacore and Guppy are similar in terms of accuracy metrics, Guppy is an order of magnitude faster (∼1 500 000 bp/s vs ∼120 000 bp/s) due to its use of GPU acceleration (Fig 1). Despite also using GPU acceleration, Chiron was the slowest basecaller tested (∼2500 bp/s), with the INF032 test set taking more than 2.5 days to basecall. This means that Chiron would take over a month to basecall a typical MinION yield of 10 Gbp, making it impractical for anything but very small read sets. Flappie also suffered from low speed performance (∼14 000 bp/s).

### Custom model performance

Running Guppy v2.2.3 with our custom-*Kp* model (trained on unamplified DNA from 30 *K. pneumoniae*, 10 other Enterobacteriaceae and 10 other Proteobacteria, see Methods and Fig S1) produced a modest increase in read accuracy (Q9.5) and a large increase in consensus accuracy (Q28.5) for the benchmarking set, relative to the default model (Fig 1). This demonstrates that there is a benefit for using taxon-specific training data. The default and custom-*Kp* models performed similarly in terms of speed.

Our custom-*Kp*-big-net model (trained on the same data as custom-*Kp* but with a larger neural network, Fig S2) delivered even further improvements in both read accuracy (Q10.4) and consensus accuracy (Q31.6), showing that more complex neural networks also have the potential to give improved results, but at a cost to speed performance. The custom-*Kp*-big-net model could not be run on the GPU because it uses neural network layers that are not pre-compiled into the Guppy program. It had to be run on the CPU instead, which along with the increased complexity of its neural network, resulted in a speed of ∼13 000 bp/s – two orders of magnitude slower than Guppy run with the default or custom-*Kp* models on GPU.

Our custom-trained models were designed for *K. pneumoniae* and performed well on the *K. pneumoniae* benchmarking set. To see if these results generalise to other genomes (both *K. pneumoniae* and more distantly-related species), we also ran all available Guppy models on additional read sets (Fig 2). The flip-flop model performed better than the default model for all genomes, with a mean improvement of +0.71 in the read qscore and +0.36 in the consensus qscore. The custom-*Kp* model performed much better than the default model for genomes in Enterobacteriaceae (*K. pneumoniae* and *S. sonnei*), with mean qscore improvements of +0.63 (read) and +4.72 (consensus). However, these benefits were not seen for species outside of Enterobacteriaceae, where the mean qscore changes were 0.00 (read) and −1.64 (consensus). This taxon-specific improvement is likely due to the custom-*Kp* model’s ability to more accurately call Dcm-methylation motifs which are found in Enterobacteriaceae7 (see details below). The improved performance of custom-*Kp*-big-net over the custom-*Kp* model was not taxon-dependent and showed mean qscore improvements of +1.01 (read) and +3.15 (consensus) across all genomes. In almost all cases, the custom-*Kp*-big-net model produced the most accurate reads and consensus, the exception being *S. aureus* where the flip-flop model produced the most accurate reads.

**Fig 2.**
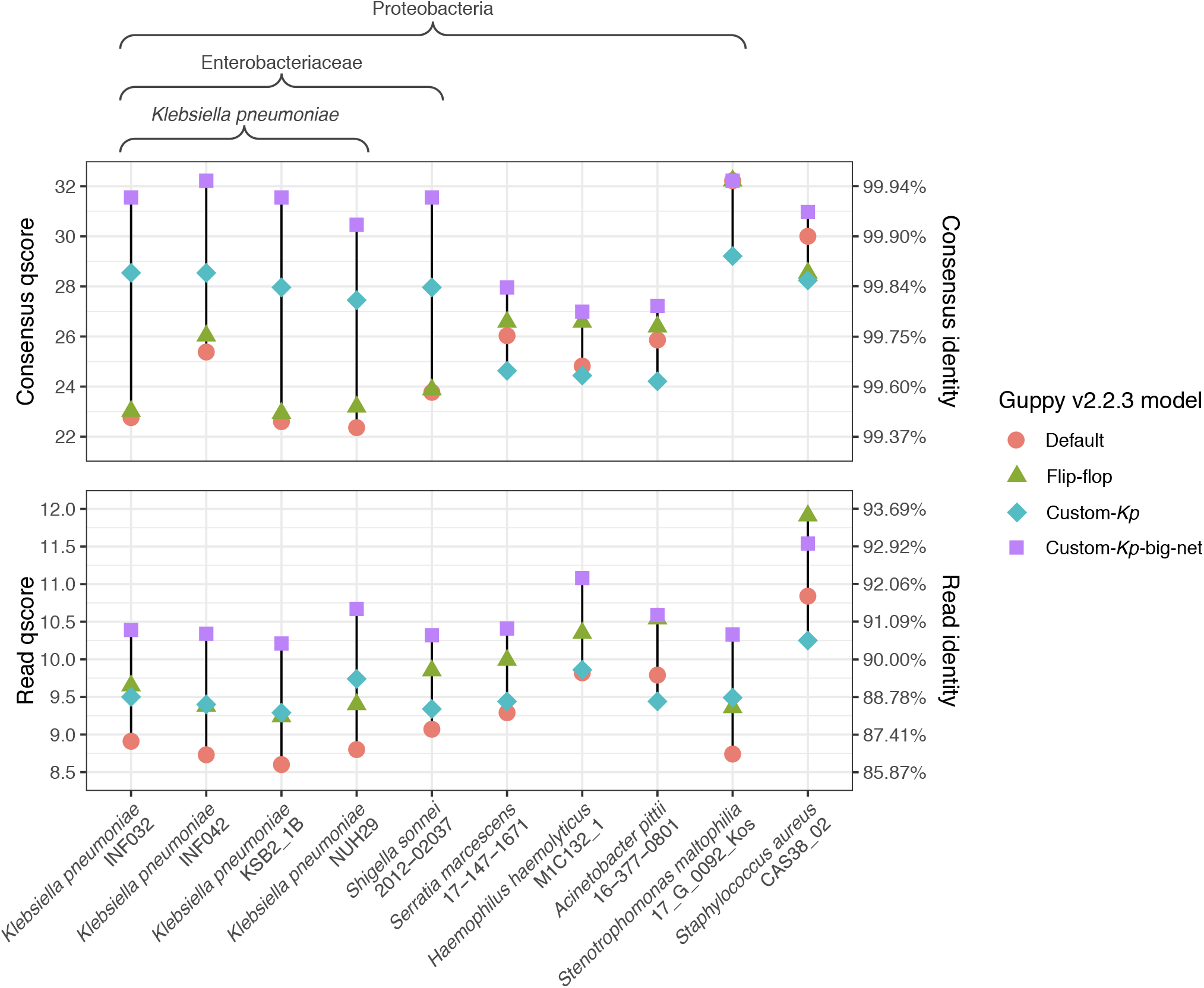
Read and consensus accuracy from Guppy v2.2.3 for a variety of genomes using different models: the default RGRGR model, the included flip-flop model and the two custom models we trained for this study. Both custom models used the same training set which focused primarily on *K. pneumoniae*, secondarily on the Enterobacteriaceae family and lastly on the Proteobacteria phylum.

Neither the custom-*Kp* model nor the custom-*Kp*-big-net model use the new neural network architecture present in Guppy’s flip-flop model. Presumably, a model based on this flip-flop architecture and trained on our custom training data would enjoy both the benefits of the flip-flop model (improved accuracy for all genomes) and of the custom-*Kp* model (improved accuracy for Enterobacteriaceae). However, the current version of Sloika (v2.1.0) does not allow for custom training with the flip-flop architecture.

### Consensus error profiles

In order to understand the impact of the various basecallers on different kinds of consensus basecalling errors, we quantified error profiles for the *K. pneumoniae* benchmarking genome in terms of the number of errors in Dcm-methylation sites, homopolymers and other sites (Fig 3). All ONT basecallers performed poorly with Dcm-methylation sites when using the default models, and these made up a large proportion of consensus errors: for current versions of ONT basecallers ∼70% of the errors were in Dcm motifs and they created ∼0.4% error relative to the reference. This implies that the models were trained on data lacking Dcm methylation and have therefore not learned to call the sites reliably. Conversely, running Guppy v2.2.3 with our custom-trained models resulted in almost no Dcm errors (∼0.002%) because Dcm methylation was well represented in our training set. The model included in versions 0.2 and 0.3 of Chiron was trained on *E. coli* reads^3^ where Dcm modifications are expected^7^, and those versions accordingly yielded very few Dcm errors (<0.025%). The model in Chiron v0.4.2 was trained on human reads and thus yielded 0.29% Dcm errors (and more errors in general).

**Fig 3.**
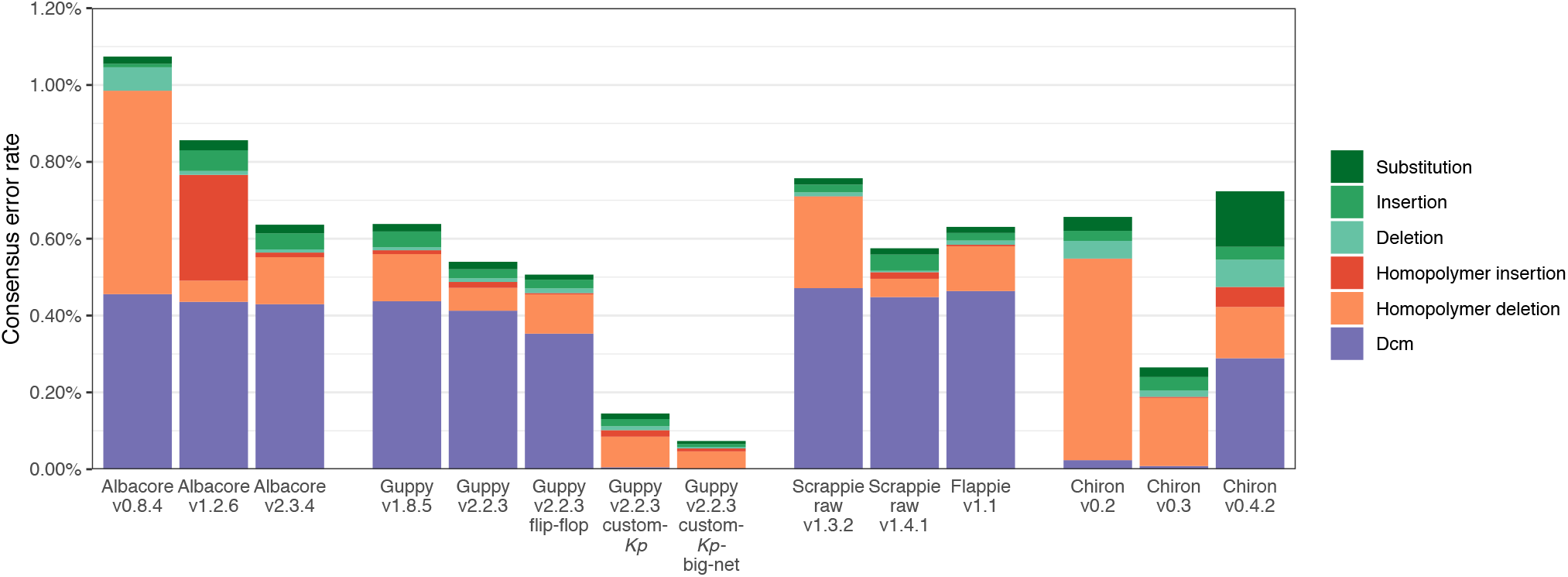
Consensus errors per basecaller for the *K. pneumoniae* benchmarking set, broken down by type. Dcm refers to errors occurring in the CCAGG/CCTGG Dcm motif. Homopolymer errors are changes in the length of a homopolymer three or more bases in length (in the reference). This plot is limited to basecallers/versions with less than 1.2% consensus error and excludes redundant results from similar versions.

After Dcm motifs, incorrect homopolymer lengths made up the majority of errors (Fig 3). ONT’s progress on this front is evident in the performance of Albacore, for which consensus accuracy improvements over time have mostly come from a reduction in homopolymer errors, from 0.53% in v0.8.4 down to 0.13% in v2.3.4. More recently, Guppy v2.2.3 has shown further improvement, bringing homopolymer errors down to 0.07%. While our custom-*Kp* model performed slightly worse than Guppy’s default model for homopolymers (0.10%), the custom-*Kp*-big-net model performed better (0.05%).

### Nanopolish performance

Nanopolish can use the raw signal data to fix errors in a consensus sequence, and it includes special logic for both Dcm methylation and homopolymers. We found that Nanopolish improved the consensus accuracy for our benchmarking set in nearly all cases, with the exception of our custom-*Kp*-big-net model (Fig 4). The pre-Nanopolish consensus accuracy was correlated with the post-Nanopolish accuracy *R*^2^=0.580), indicating that a basecaller’s consensus accuracy still matters even if Nanopolish is also used. While Nanopolish was able to account for Dcm methylation (using its `--methylation-aware` option), it often only corrected ∼70–80% of Dcm errors (Fig S4). Accordingly, the rate of Dcm errors in the pre-Nanopolish assembly was the strongest predictor of post-Nanopolish accuracy *R*^2^=0.809), with the best results (>Q29 consensus) coming from the four basecallers with very low Dcm error rates (Chiron v0.2–v0.3 and both custom models). The effect of additional rounds of Nanopolish was tested on the Guppy v2.2.3 assembly and gave only a small increase in accuracy (from Q27.5 after one round to Q28.3 after four rounds, Fig S5).

**Fig 4.**
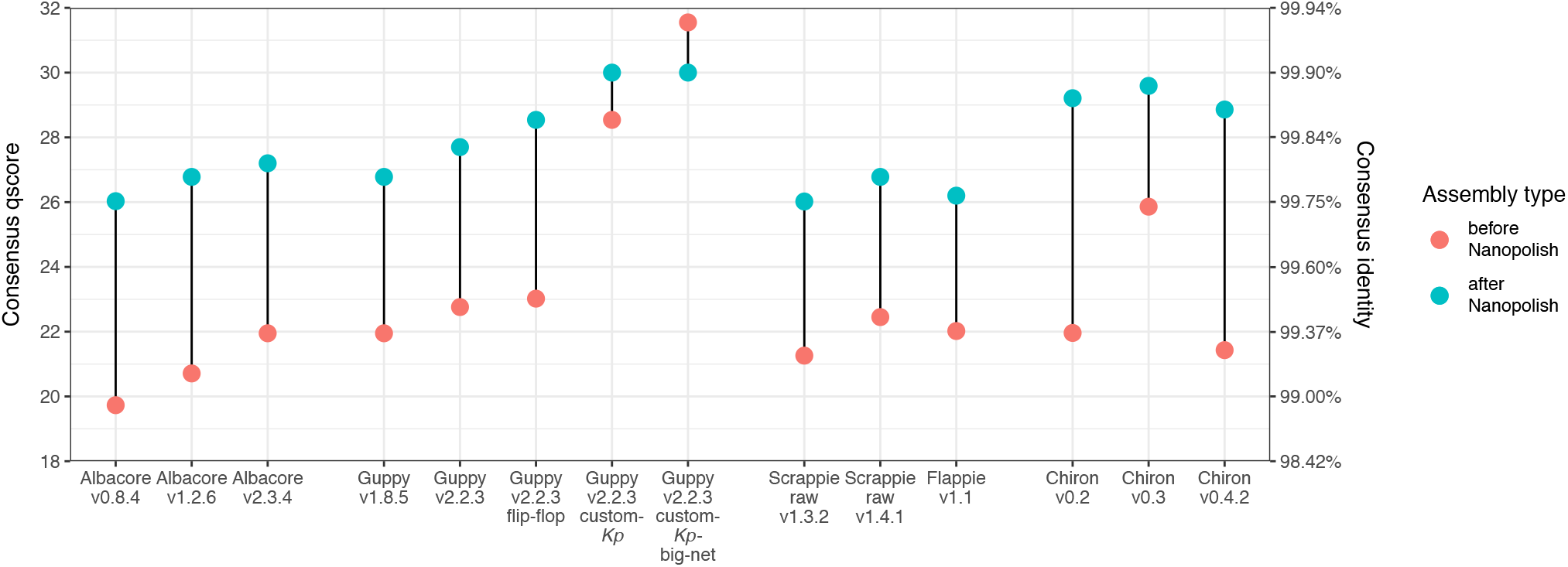
Consensus accuracy before (red) and after Nanopolish (blue) for the assemblies of *K. pneumoniae* benchmarking set.

DNA base substitutions (often referred to as single nucleotide polymorphisms or SNPs) identified from pathogen whole-genome sequence comparisons are now used routinely by public health and infection control laboratories to investigate suspected outbreaks of foodborne and other infectious diseases^8^. ONT platforms could potentially be useful in such investigations due to their portability and cost effectiveness for small sample sizes. However, the smallest number of substitution errors we encountered in a genome assembly (using Guppy v2.2.3 with the custom-*Kp*-big-net model and Nanopolish) was 337 substitutions (Fig S6, File S1), which at ∼10 times the number of true SNPs expected between bacterial pathogen genomes linked to the same outbreak8 would constitute an unacceptably high false positive rate for this application. While it is possible that tailored strategies for SNP calling from ONT reads could reduce the number of false positives, assembly-based sequence comparisons (which are frequently used in public health labs^9, 10^) would require dramatic improvements in basecalling accuracy.

## Conclusions

Best results, in terms of both read and consensus accuracy for the *K. pneumoniae* benchmarking set, were obtained using Guppy v2.2.3 with a custom model trained on data mostly from the same species (Fig 1). This superior performance seems to largely come from correct handling of Dcm methylation (Fig 3). Since DNA modification patterns can differ between taxa, we propose our results may represent a more general trend: native DNA basecalling accuracy is best when the model was trained on native DNA from the same species or a sufficiently close relative to have similar DNA modifications. For example, a model trained on native human DNA may also perform well on native mouse DNA and vice versa, as CpG methylation is common to both species^11^. The benefits of custom training may further extend to direct RNA sequencing, where base modifications can be more extensive than in DNA^12, 13^.

For most basecallers, full information is not disclosed on the taxa and DNA type (native or amplified) used to train the default model. We encourage developers to be more transparent in this regard and to consider providing multiple trained models when possible (e.g. amplified, native human, native *E. coli*, etc.) so users can choose one which most closely matches their organism. For users with sufficient quantities of training data, high-performance computers with GPUs and computational expertise, we recommend custom model training to maximise basecalling accuracy. Our custom-*Kp*-big-net model shows that even more accurate results are possible with bigger neural networks, including substantial improvements in read-level accuracy, but at a cost to speed performance (Fig 1).

ONT sequencing has seen enormous gains in both yield and accuracy over the past few years, but our results show there is still much room for improvement. Across all basecallers, models and genomes, the best consensus accuracy we observed was Q32.2 (99.94% identity). This equates, on average, to ∼3000 errors in a 5 Mbp genome. Many of these errors are substitutions which could lead to false-positive SNP calls, a potentially major impediment to outbreak investigations. In order to achieve a perfect bacterial genome assembly, the consensus accuracy will need to be orders of magnitude higher, e.g. Q70 (one error per 10 Mbp). Progress will likely come from many fronts: changes in technology and chemistry, improvements in basecalling, and development of post-assembly polishing tools. Until this goal is reached, hybrid assembly or polishing with Illumina reads will remain a necessity for researchers that depend on highly accurate sequences.

## Methods

### Custom model training

Sloika (github.com/nanoporetech/sloika) is ONT’s neural network training toolkit which can be used to make models for use in Guppy. To explore the effect of the training set on basecalling performance, we used Sloika v2.1 to train a model (‘custom-*Kp*’) tailored to *K. pneumoniae*. The training reads came from 50 different isolate genomes: 30 *K. pneumoniae* (chosen based on their phylogenetic uniqueness, each from a different lineage), 10 from other species of Enterobacteriaceae and 10 from other families of Proteobacteria (Fig S1, File S1). Our training reads came from 20 different MinION runs, 10 of which were barcoded runs that contributed multiple genomes to the training set. Illumina reads were also available for all genomes, and we used SKESA14 (v2.3.0) to produce high-quality contigs for each.

From an initial collection of 5 629 714 reads, we trimmed each read’s signal at the fast5 level, removing low-variance open-pore signal5 and then an additional 2000 signal values from the start and end of the reads which served to remove adapter and barcode signals. We also discarded short reads (<50 000 signal values), leaving 1 985 997 trimmed reads. We basecalled these reads (using Albacore v2.3.4) to select reads based on length (>5 kbp), completeness of alignment to the SKESA contigs (<30 bp unaligned) and quality over a sliding window (no indel regions exceeding 25 bp in size). This further reduced our set to 766 551 reads. We filtered the reads once more (following Sloika’s training instructions: `example_training.sh`), this time on the quality of the raw-signal-to-reference alignment. We set aside 20 of the resulting reads from each genome (1000 in total) to use as a validation set, leaving 226 166 for training the neural network. Sloika subdivides these reads into ‘chunks’ of 4000 signal values, of which there were 7 693 885.

Using Sloika with this entire training set would have required hundreds of gigabytes of RAM, so we produced a fork of Sloika (github.com/rrwick/sloika) modified to load a random subset of the training data at periodic intervals. By only training on 5% of the data at a time and reloading a new 5% every 250 training batches, we were able to use the entirety of our training data while keeping RAM usage under 10 GB. We trained the custom-*Kp* model for 47 500 batches (100 chunks per batch) on an NVIDIA P100 GPU, which took 36.5 hours.

Albacore, Guppy and Scrappie all use an architecture that ONT calls RGRGR – named after its alternating reverse-GRU and GRU layers (Fig S2, top-left). To test whether more complex networks perform better, we modified ONT’s RGRGR network by widening the convolutional layer and doubling the hidden layer size (Fig S2, top-right). We trained this ‘custom-*Kp*-big-net’ model on the same bacterial training set using our fork of Sloika. Training on an NVIDIA P100 GPU for 44 000 batches took 48 hours.

### Read sets

To test basecaller performance, we used a set of reads generated using a MinION R9.4 flowcell to sequence native DNA extracted from the bacterium *Klebsiella pneumoniae*. The bacterial sample (isolate INF032, BioSample accession SAMEA3356991) was isolated from a urinary tract infection in an Australian hospital^15^. It was sequenced as part of a barcoded MinION run with other *Klebsiella* isolates, following the DNA extraction and library preparation protocol described in Wick et al. 2017^16^. This particular sample was chosen for benchmarking basecalling accuracy because it had a good yield of ONT reads (see below) and contained no plasmids, making for a simpler assembly. It is not in the same *K. pneumoniae* lineage as any of the genomes used to train our custom models (Fig S1). Since the sequenced DNA was native, it contains base modifications, the most relevant of which is Dcm methylation (conversion of cytosine to 5-methylcytosine at particular motifs) which is common in some species of Enterobacteriaceae^7^. High-quality Illumina reads were available for this sample: DNA was extracted and sequenced as 125 bp paired-end reads via Illumina HiSeq 2000 at the Sanger Institute, producing 5 455 870 reads (ENA accession ERR1023765) with 133× read depth over the INF032 chromosome^15^. This allowed us to generate an accurate reference sequence via hybrid assembly using Unicycler16 (v0.4.0) which produced a single 5 111 537 bp contig with a GC-content of 57.6%. This reference was used as the ground truth sequence when assessing read and consensus accuracy for each basecaller (see below).

A subset of ONT reads was extracted for benchmarking basecallers against the reference genome of INF032. The entire barcoded MinION run containing INF032 was demultiplexed using Deepbinner^17^, using its `--require_both` option for high-precision demultiplexing. This produced 70494 reads (∼1.1 Gbp) for the barcode corresponding to INF032. We further reduced this dataset by basecalling with Guppy v1.6.0 (the current version at the time of read selection), aligning the resulting reads (using minimap218 v2.14) to the INF032 reference genome and selecting those with a ≥22 kbp alignment to the reference. This served to exclude ‘junk’ reads, very low-quality reads, improperly demultiplexed reads (belonging to a different isolate) and short reads. The threshold of ≥22 kbp was chosen because it reduced the dataset to approximately 100× mean read depth for the INF032 genome while maintaining even coverage (Fig S3). The resulting set contained 15 154 reads with lengths ranging from 22–134kbp (N50=37 kbp) and totalling ∼550Mbp. These reads are significantly longer than the longest repeat in the INF032 genome (the ∼5.5 kbp rRNA operon) so each can be reliably mapped to its correct location on the reference genome. The Guppy qscore distributions (Fig S3) show that while the selection process removed the lowest quality reads, the resulting reads still span a wide quality range, with 458 (∼3%) falling below ONT’s ‘fail’ threshold of Q7. This read set (hereafter referred to as the ‘benchmarking set’) was used with all basecallers and versions.

In addition to the primary *K. pneumoniae* benchmarking set, we also prepared test read sets for nine additional genomes. These include three other *K. pneumoniae* genomes and one genome each from six different bacterial species: *Shigella sonnei, Serratia marcescens, Haemophilus haemolyticus, Acinetobacter pittii, Stenotrophomonas maltophilia* and *Staphylococcus aureus*. As well as covering a wider range of species, these read sets span a wider date range (Feb 2017–Aug 2018) and include both R9.4 and R9.4.1 flowcells (File S1). They were prepared in the same manner as the benchmarking set, but we adjusted the alignment length threshold for each set as appropriate for the genome and read depth (3 kbp to 33 kbp, see File S1). These read sets (hereafter referred to as the ‘additional sets’) were used to more thoroughly assess the current version of Guppy (v2.2.3) using its default model, its included flip-flop model and our two custom models (custom-*Kp* and custom-*Kp*-big-net).

### Read and consensus accuracy

We ran all basecallers on the benchmarking read set, with each producing either a FASTQ or FASTA file suitable for downstream analysis. To allow a comparison of speed performance, all basecalling was carried out on the same computer: six core (12 thread) Intel Xeon W-2135 CPU, 32 GB RAM, NVIDIA GTX 1080 GPU and 1 TB NVMe SSD. Basecalling was carried out using all 12 CPU threads, or if supported by the basecaller, on the GPU.

To assess read accuracy, we aligned each basecalled read set to the reference INF032 genome using minimap218 (v2.12). Each read’s identity was defined as the number of matching bases in its alignment divided by the total alignment length including insertions and deletions, a.k.a. the ‘BLAST identity’ (lh3.github.io/2018/11/25/on-the-definition-of-sequence-identity). If less than half of a read aligned to the reference, it was deemed unaligned and given an identity of 0%. Read accuracy distributions had a considerable amount of variance, ranging from ∼95% identity for the best reads down to ∼60%, below which reads were too noisy to be aligned by minimap2. We therefore defined each basecaller’s overall read accuracy as the median identity of the reads in the set.

We used Rebaler (github.com/rrwick/Rebaler) to generate a consensus sequence from each basecalled read set. Rebaler is a reference-based assembler written for the purpose of comparing basecallers. It works by first replacing all parts of the reference genome using read sequences and then polishing the genome with multiple rounds of Racon^19^. This approach ensures that the assembled genome will have the same large-scale structure as the reference, but small-scale details (e.g. basecalls) will not be affected by the reference sequence. Even after multiple rounds, Racon does not always converge to the best possible sequence, so we used Rebaler with an iterative approach: running the assembly multiple times, each with shuffled input reads and a rotated (shifted start position) reference genome. We used 10 iterations, each resulting in a slightly different assembly which were then used as the ‘reads’ for a final Rebaler assembly. This iterative approach was able to reduce assembly errors by about 20%: individual Rebaler assemblies of the Guppy v2.2.3 reads had a consensus accuracy of 99.33% (Q21.74) but the final iterative assembly had an accuracy of 99.47% (Q22.76).

To assess consensus accuracy, we divided each final Rebaler assembly into 10 kbp pieces and analysed them in the same manner used for the reads: aligning to the reference and calculating identity. Each basecaller’s overall consensus accuracy was defined as the median identity of these 10 kbp pieces. The distributions of identities for assembly pieces were narrow and had standard deviations of <0.1%.

To classify consensus sequence errors by type, we aligned each assembly to the reference using NUCmer20 (v3.1) and then classified each error based on the reference context. An error was classified as ‘Dcm’ if it occurred in a Dcm-methylation motif (CCAGG or CCTGG). It was classified as ‘homopolymer insertion’ or ‘homopolymer deletion’ if the error added or removed a base from a homopolymer three or more bases in length. If the previous categories did not apply, the error was classified as ‘insertion’, ‘deletion’ or ‘substitution’ as appropriate.

### Polishing

This study is focused on the performance of basecalling tools, and post-assembly polishing with raw signal data is a separate topic that falls outside our scope. However, many users who produce ONT-only assemblies will run Nanopolish on their result, which uses the raw read signals to improve the consensus accuracy of an assembly. This raises the question: does basecaller choice matter if Nanopolish is used downstream? To assess this, we ran Nanopolish v0.10.2^21^ on each final Rebaler assembly and assessed the consensus accuracy as described above.

## Acknowledgments

We would like to thank ONT staff for answering questions about commands and resource requirements to run the Sloika software to train custom models and the commands needed to deploy these models in Guppy.

## Data availability

All read sets (raw fast5 and basecalled), assemblies, reference genomes, training data and custom models used in this paper are available via figshare (10.26180/5c5a5f5ff20ed, 10.26180/5c5a5fa08bbee, 10.26180/5c5a5fb6ac10f, 10.26180/5c5a5fc61e7fa, 10.26180/5c5a5fcf72e40, 10.26180/5c5a7292227de).

Scripts used to perform basecalling and analysis are available via GitHub: https://github.com/rrwick/Basecalling-comparison (10.5281/zenodo.1043611).

## Supplementary material

**File S1. supplementary_tables.xlsx**

This spreadsheet contains information on each test read set, basecaller commands, all accuracy and error data results, and information on the training sets used for the custom models.

**Fig S1.**
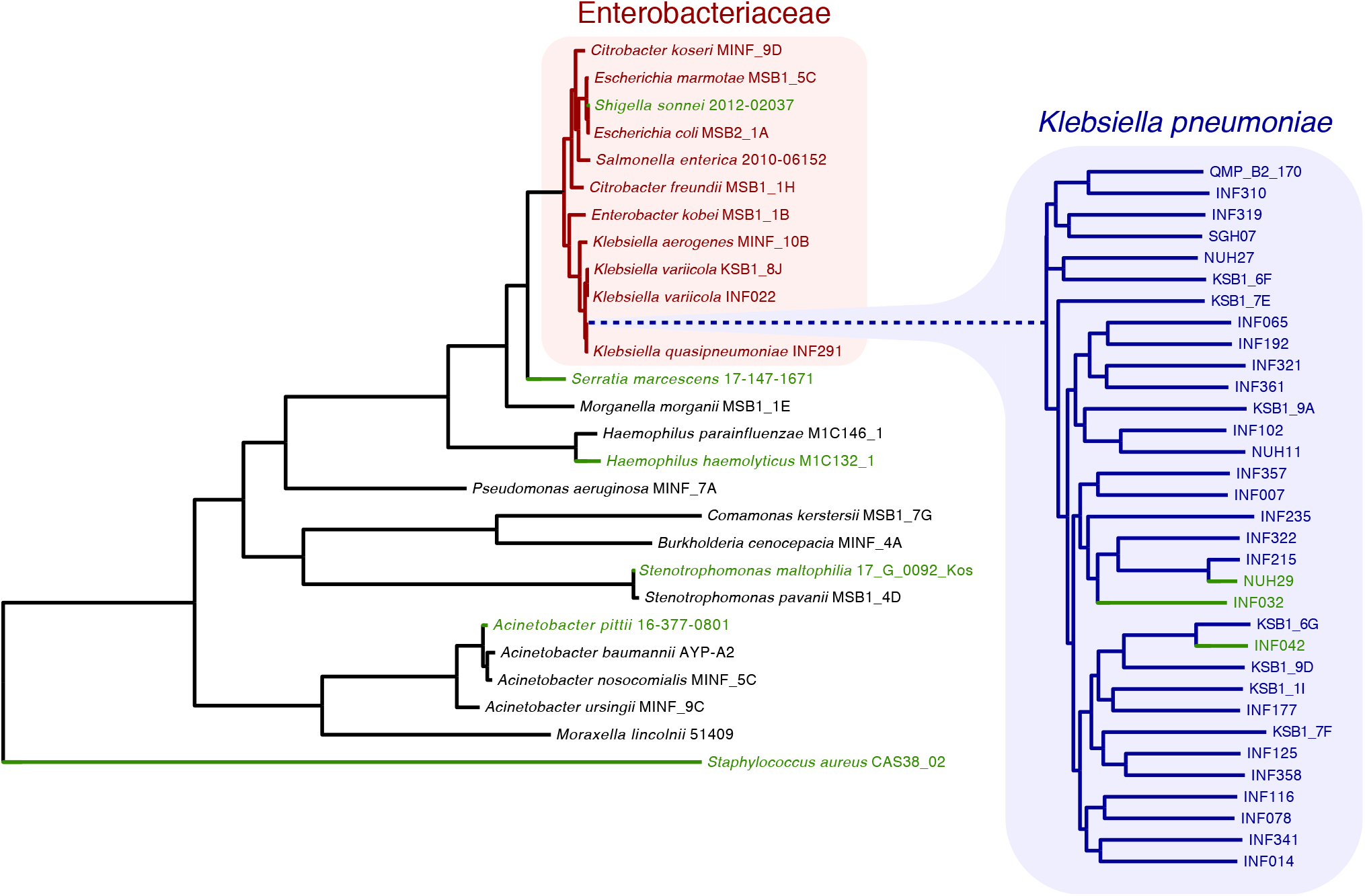
Tree of the genomes used for custom Sloika training. All isolates are from the Proteobacteria phylum, with an emphasis on the Enterobacteriaceae family (red) and *Klebsiella pneumoniae* (blue). The genomes used to test base-callers/models are also included in this tree in green, but were not used to make training data.

**Fig S2.**
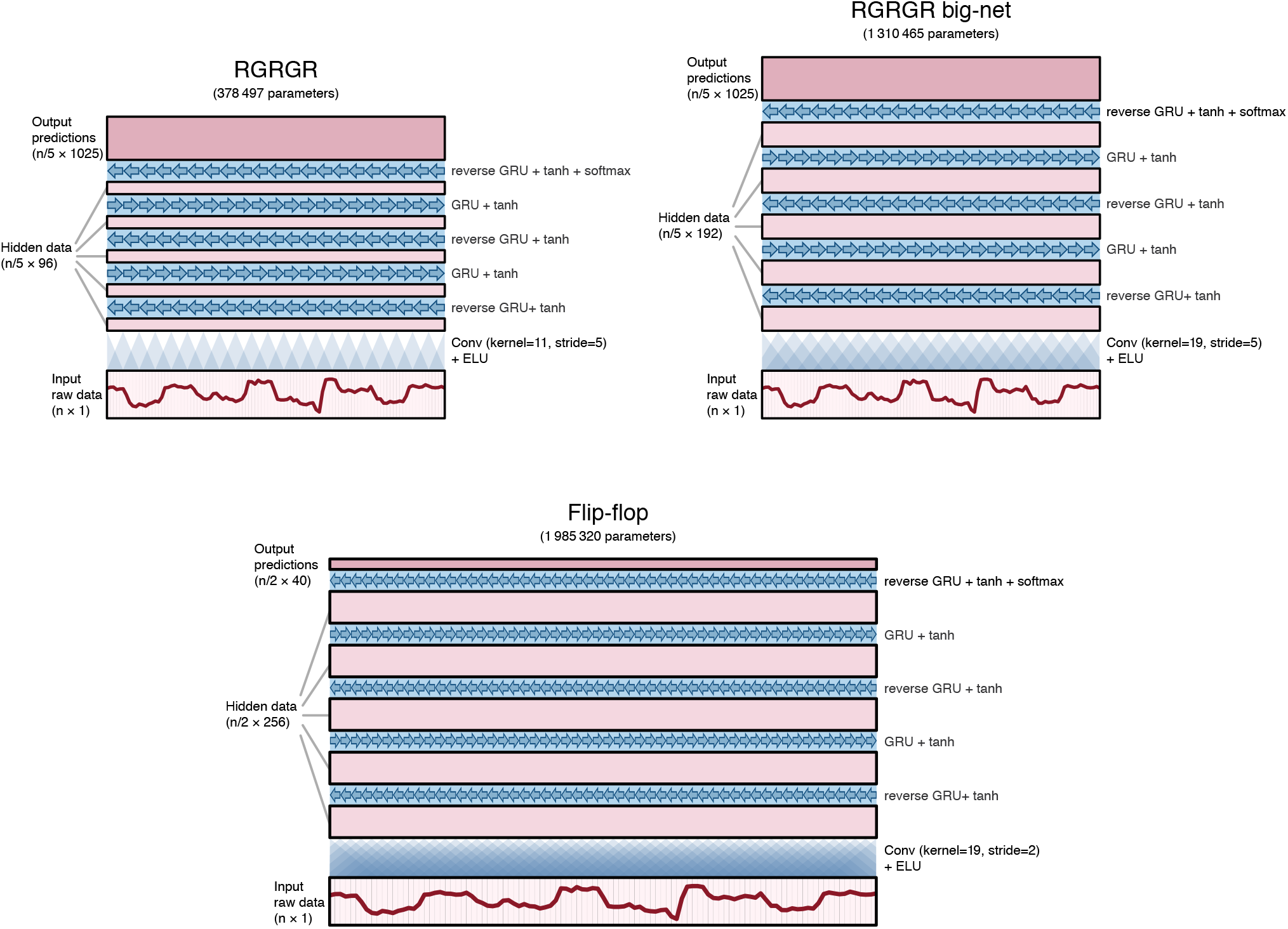
ONT neural network architectures. The RGRGR network (left) is used in Albacore, Guppy and Scrappie raw, and it was used to train the custom-*Kp* model. The big-net (right) is a modification of the RGRGR network, with a wider convolution and larger hidden data. For both networks, the convolutional layer reduces the data length (n) by a factor of five (i.e. there are five raw data values per predicted state). There are 1025 possible predicted values: 1024 for each possible 5-mer plus an additional ‘stay’ state.

**Fig S3.**
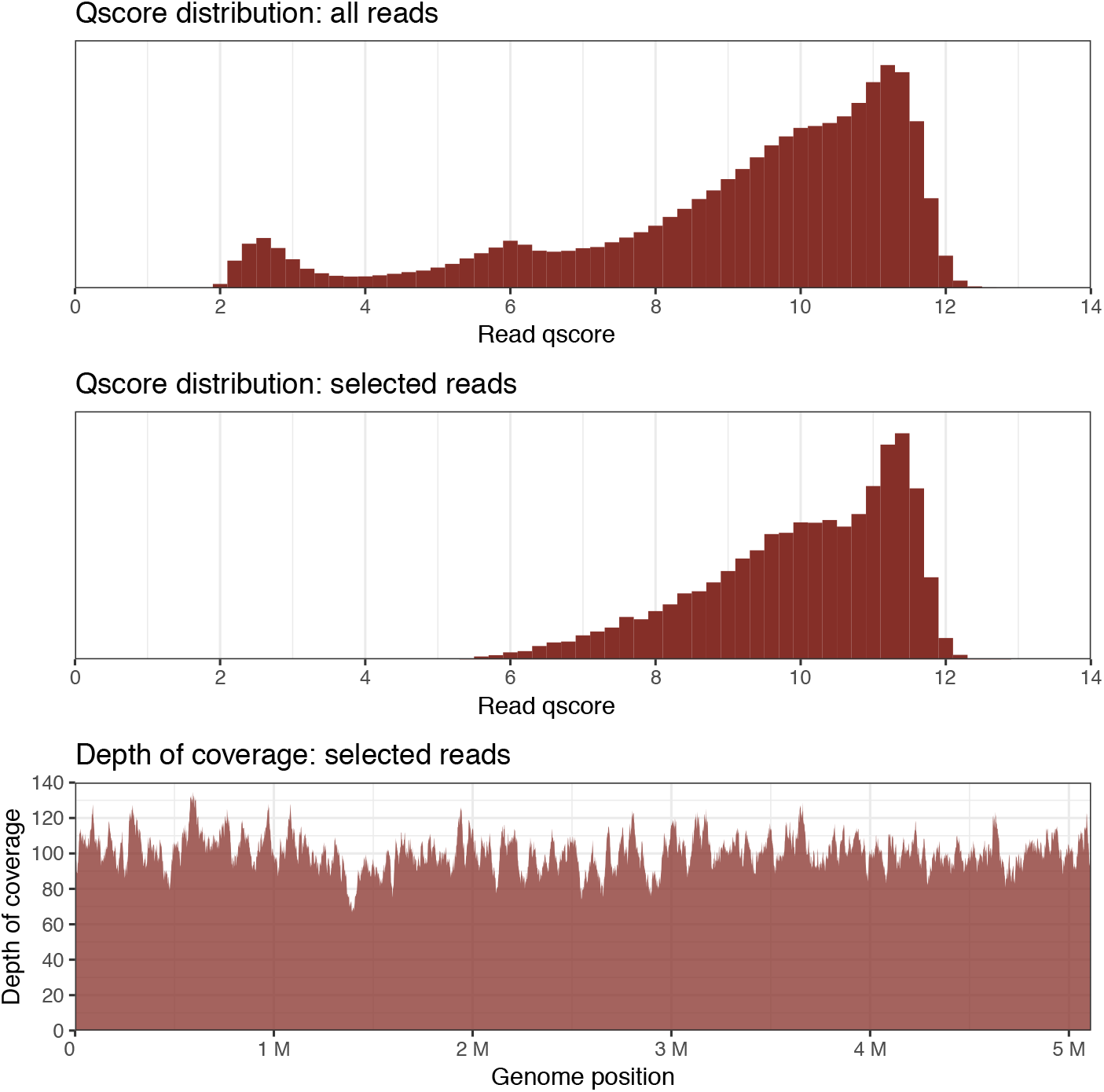
Qscore distributions (according to Guppy v1.6.0) for the entire barcoded MinION run (top) and the selected INF032 reads (bottom).

**Fig S4.**
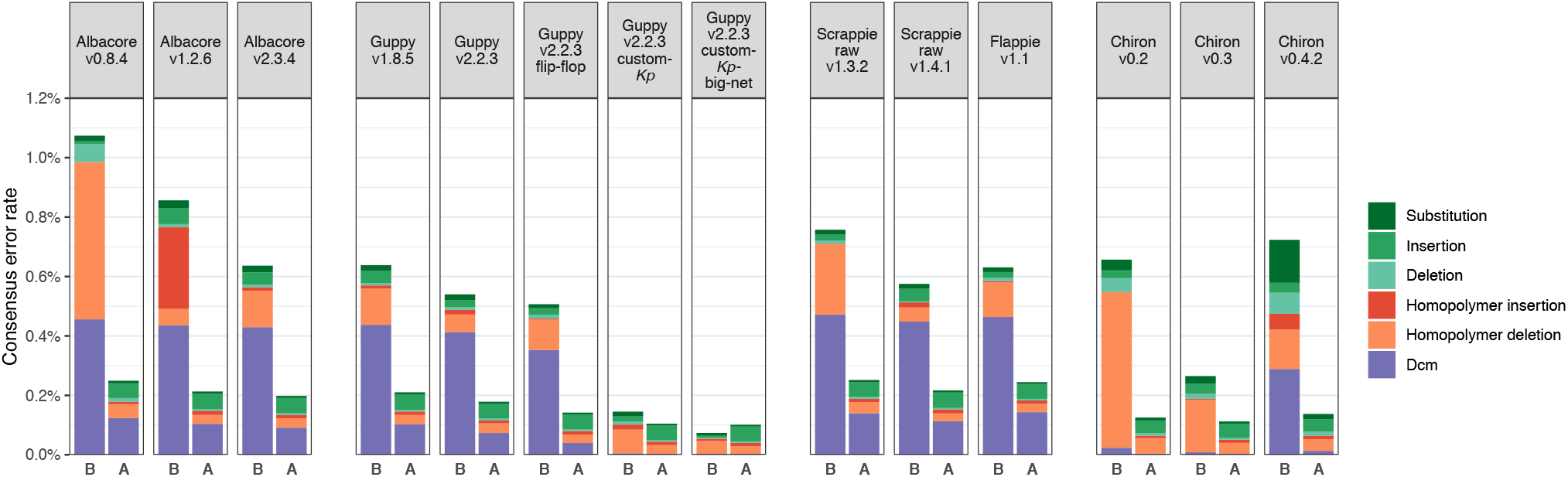
Consensus errors per basecaller, broken down by type (same types as Fig 3), before (B) and after (A) Nanopolish. Non-Dcm errors are relatively occur at a similar rate in each post-Nanopolish assembly, but the number of Dcm errors depends on their prevalence in the pre-Nanopolish assembly.

**Fig S5.**
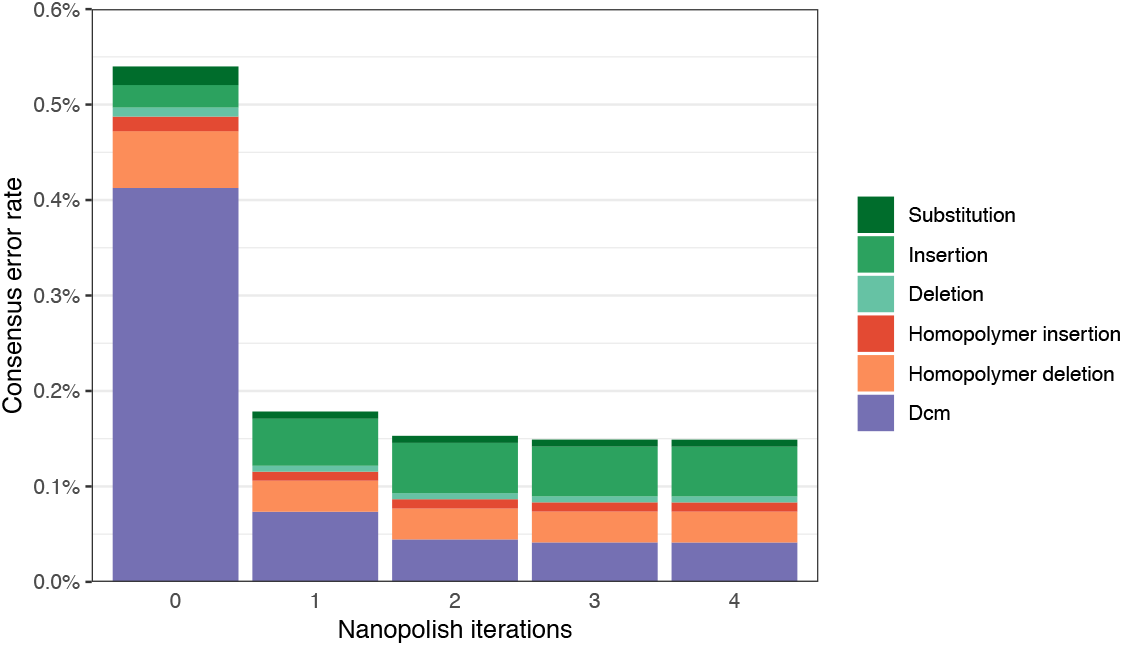
Consensus errors with repeated iterations of Nanopolish, using the Guppy v2.2.3 dataset. Accuracy improves only slightly with successive iterations, with most of the improvement coming from Dcm errors.

**Fig S6.**
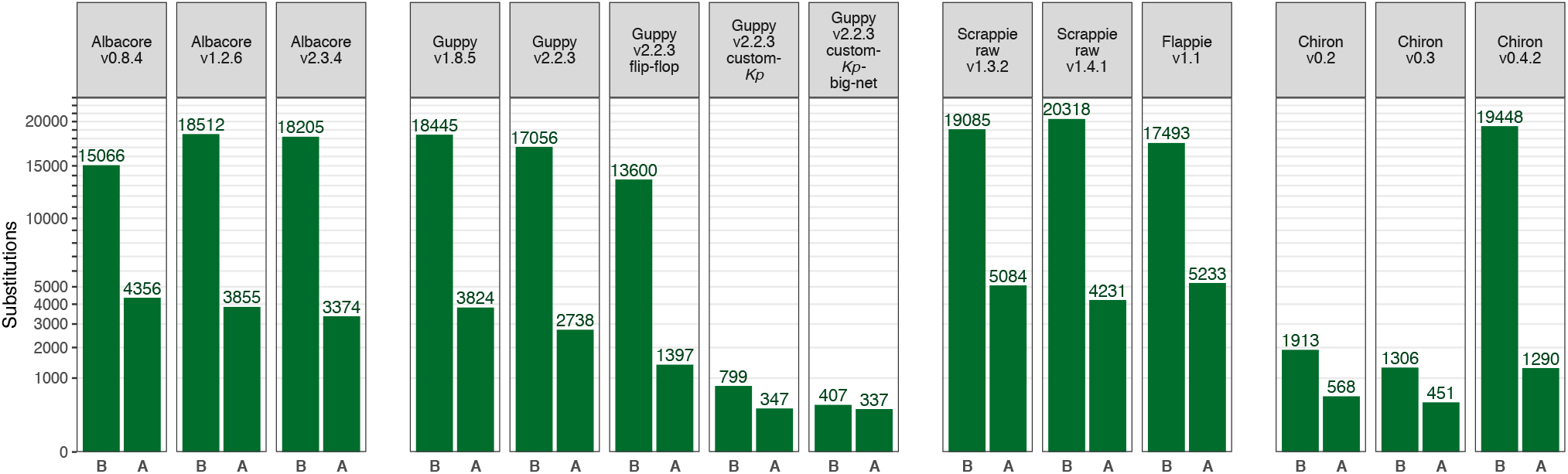
Number of consensus sequence substitution errors per basecaller, before (B) and after (A) Nanopolish. All substitution types (both Dcm-related and non-Dcm-related) are counted.

